# Extra base hits: widespread empirical support for instantaneous multiple-nucleotide changes

**DOI:** 10.1101/2020.05.13.091652

**Authors:** Alexander G Lucaci, Sadie R Wisotsky, Stephen D. Shank, Steven Weaver, Sergei L. Kosakovsky Pond

**Affiliations:** Institute for Genomics and Evolutionary Medicine, Temple University, Philadelphia, PA, USA

**Keywords:** multiple nucleotide substitutions, positive selection, polymerase zeta, evolutionary process, adaptive evolution

## Abstract

Despite many attempts to introduce evolutionary models that permit substitutions that instantly alter more than one nucleotide in a codon, the prevailing wisdom remains that such changes are rare and generally negligible (or are reflective of non-biological artifacts, such as alignment errors), and codon models continue to posit that only single nucleotide change have non-zero rates. We develop and test a simple hierarchy of codon-substitution models with non-zero evolutionary rates for only one-nucleotide (1H), one- and two-nucleotide (2H), or any (3H) codon substitutions. Using 35,000 empirical alignments, we find widespread statistical support for multiple hits: 58% of alignments prefer models with 2H allowed, and 22% – with 3H allowed. Analyses of simulated data suggest that these results are not likely to be due to simple artifacts such as model misclassification or alignment errors. Further modeling revealed that synonymous codon island jumping among codons encoding serine, especially along short branches, contributes significantly to this 3H signal. While serine codons were prominently involved in multiple-hit substitutions, there were other common exchanges contributing to better model fit. It appears that a small subset of sites in most alignments have unusual evolutionary dynamics not well explained by existing model formalisms, and that commonly estimated quantities, such as dN/dS ratios may be biased by model misspecification. Our findings highlight the need for continued evaluation of assumptions underlying workhorse evolutionary models and subsequent evolutionary inference techniques. We provide a software implementation for evolutionary biologists to assess the potential impact of extra base hits in their data in the HyPhy package.

## Introduction

Most modern codon models in wide-spread use assume any changes within a codon happen as a sequence of single instantaneous nucleotide change, enforced by setting instantaneous rates between codons that differ in more than one nucleotides to zero. This choice was made independently for the mechanistic models of Muse and Gaut (1994) and Goldman and Yang (1994), and adopted by subsequent model developers and practitioners. For example, when Halpern and Bruno (1998) introduced their mutation-selection models, they considered the general multi-hit (MH) case first, but noted that introducing the single hit assumption “..has very little effect on our results under the conditions we have investigated.” This assumption is both computationally convenient and biologically sound in the majority of cases, since it assumes that the events when randomly occurring mutations occur instantaneously in the same codon is vanishingly rare. While rare, evidence for substitutions occurring in tandem at adjacent nucleotide sites had been reported at about the same time the codon models were being introduced (Wolfe and Sharp, 1993). Averof *et al*. (2000) reported significant rates of changes between TCN and AGY codon islands in perfectly conserved serine residues, and argued against going through intermediary non-synonymous changes due to their likely deleterious effects, while (Rogozin *et al*., 2016) argued that strong purifying selection on single nucleotide changes is a more plausible explanation in general. Neither of those studies has considered an explicit evolutionary model, however. Serine is the only amino-acid with synonymous codon islands in the universal genetic code, but several other codes have have other aminoacids with this property: leucine in the *Chlorophycean* and *Scenedesmus obliquus* mitochondrial codes (TAG and CTH), and alanine in the *Pachysolen tannophilus* nuclear code (CTG and GCH).

Recent studies estimate that 2% of nucleotide substitutions are part of larger multiple nucleotide changes that occur simultaneously (Harris and Nielsen, 2014; Kaplanis *et al*., 2019), due in part to an error-prone DNA polymerase Zeta. Human germline tandem mutations have been estimated to constitute 0.4% of all mutations (Chen *et al*., 2014), and individual cases of such mutations have been reported to have significant phenotypic consequences, e.g. via their effects on protein folding (Okada *et al*., 2017).

A number of codon model extensions have incorporated MH, invariably finding improvement in fit and (if the model allowed testing) statistically significant evidence of non-zero rates involving multiple nucleotide changes. Kosiol *et al*. (2007) developed a general MH empirical codon substitution model estimated jointly from a large collection of training alignments, and noted that it was overwhelmingly preferred to standard SH models on a sample of biological data from the Pandit database. Several groups have independently developed alternative codon model parametrizations to allow for MH, including Whelan and Goldman (2004) (“… these events [MH] are far more prevalent than previously thought”), Zaheri *et al*. (2014), and Dunn *et al*. (2019) (the latter two studies show a dramatically better model fit to empirical alignments when allowing MH). Other studies that used evolutionary models with some support for MH based on, at least in part, numerical rate estimates from training data include De Maio *et al*. (2013); Doron-Faigenboim and Pupko (2007); Miyazawa (2011); Zoller and Schneider (2012) Despite multiple introductions to the field, these models have not been able to gain a substantial foothold in applied evolutionary analyses, and for some of these methods, software implementing them is no longer available.

Failure to include multiple hits in codon substitution models may mislead evolutionary hypothesis testing. Venkat *et al*. (2018) found that the addition of a double-hit rate parameter improved model fit and impacted branch-specific inferences of positive selection (MH along short branches can inflate false positives). Dunn *et al*. (2019) used principled simulation studies to show that fitting 1H models to data generated with low rates of multiple hits can increase false positive rates and dilute power for identifying individual sites subject to positive selection.

In this study we develop simple extensions to the Muse and Gaut (1994) based codon model which add double, and triple instantaneous (2H, 3H) changes and compares them to simpler models in large collections of empirical data. Our models are mechanistic and simpler than those proposed by Whelan and Goldman (2004) and Dunn *et al*. (2019). This relative simplicity allows our models to be implemented and fitted quickly, and offers straightforward interpretation, including the ability to identify individual sites that benefit from the addition of MH. The primary goal of our data analyses is to establish how often evidence for multiple hits can be detected in large-scale empirical databases (something that no other study looking at evolutionary models has done), identify the codons that are frequently involved in such events, and explore plausible biological explanations for why these rates are non-zero for a majority of alignments.

## Results

### Benchmark alignments

We introduce the models using a collection of thirteen representative alignments that we and others have been using to benchmark selection analyses, most recently in Wisotsky *et al*. (2020). We also consider the primate lysozyme alignment originally analyzed with codon models by Yang (1998). We consider five models (see Table 1 and the methods section for details), which form a nested hierarchy (with the exception of 3HSI and 3H which are not nested), each with one additional alignment-wide parameter.

**Table 1.** Key parameters, models considered here. GDD = general discrete distribution; 1H, 2H, 3H – instantaneous changes involving one, two, or three nucleotides.

| Parameter | Description | Model |  |  |  |  |
| --- | --- | --- | --- | --- | --- | --- |
|  |  | 1H | 2H | 3HSI | 3H | 3H+ |
| $\omega_i$ | Site dN/dS ratio | Random effect 3-bin GDD distribution | | | | |
| $\delta$ | Global 2H/1H rate ratio | 0 | Estimated | Estimated | Estimated | Estimated |
| $\psi_s$ | Global 3H/1H rate ratio for synonymous codon islands | 0 | Estimated | Estimated | $=\psi$ | Estimated |
| $\psi$ | Global 3H/1H rate ratio | 0 | 0 | 0 | Estimated | Estimated |

**Table 2.** Analysis of benchmark datasets. *N* - number of sequences, *S* - number of codons, *T* - total tree length (expected subs/site) under the 1H model, *δ* rate estimate under the 3H model (2H model in parentheses), *Ψs* estimate under the 3H model, *Ψ* estimate under the 3H model. Likelihood ratio p-values for pairwise model tests, e.g. 2H:1H – 2H alternative, 1H null. Values *<* 0.05 are bolded. # sites with *ER>* 5 lists the number of sites which show strong preferences for 2H or 3H model using evidence ratios of at least 5 (see text)

| Gene | N | S | T | $\delta$ | $\psi_s$ | $\psi$ | LRT p-value | | | | | # sites with $ER > 5$ | |
| --- | --- | --- | --- | --- | --- | --- | --- | --- | --- | --- | --- | --- | --- |
|  |  |  |  |  |  |  | 2H:1H | 3H+:1H | 3H+:2H | 3H+:3HSI | 3HSI:2H | 2H:1H | 3H+:2H |
| β-globin | 17 | 144 | 2.5 | 0.7 (0.81) | $> 100$ | 0 | **<0.001** | **<0.001** | **<0.001** | 1 | **<0.001** | 10 | 6 |
| Flavivirus NS5 | 18 | 342 | 6 | 0.49 (0.73) | 2.3 | 0.6 | **<0.001** | **<0.001** | 0.056 | 0.062 | 0.13 | 16 | 0 |
| Primate Lysozyme | 19 | 130 | 0.24 | 0 (0) | 0 | 0 | 1 | 1 | 1 | 1 | 1 | 0 | 0 |
| COXI | 21 | 510 | 5.3 | 0.4 (0.4) | 0 | 0 | **<0.001** | **0.0018** | 1 | 0.94 | 0.98 | 3 | 0 |
| Drosophila *adh* | 23 | 254 | 1.4 | 0.31 (0.4) | 0 | 0.42 | **<0.001** | **<0.001** | 0.19 | 0.067 | 0.99 | 4 | 0 |
| Encephalitis *env* | 23 | 500 | 0.84 | 0.076 (0.076) | 0 | 0 | 0.19 | 0.42 | 1 | 1 | 0.98 | 0 | 0 |
| Sperm lysin | 25 | 134 | 2.8 | 0.4 (0.46) | 2.3 | 0.3 | **<0.001** | **<0.001** | **0.04** | **0.015** | 0.49 | 21 | 1 |
| HIV-1 *vif* | 29 | 192 | 0.96 | 0.007 (0.044) | 0 | 0.17 | 0.058 | **0.0013** | **0.0077** | **0.0018** | 0.95 | 0 | 2 |
| Hepatitis D virus antigen | 33 | 196 | 1.9 | 0.34 (0.37) | 0 | 0.2 | **<0.001** | **<0.001** | 0.25 | 0.098 | 0.99 | 15 | 0 |
| Vertebrate Rhodopsin | 38 | 330 | 3.9 | 0.54 (0.72) | 9.2 | 0.9 | **<0.001** | **<0.001** | **<0.001** | **<0.001** | **0.0029** | 43 | 3 |
| Camelid VHH | 212 | 96 | 15 | 0.29 (0.32) | 0 | 0.13 | **<0.001** | **<0.001** | **0.011** | **0.0026** | 0.92 | 46 | 0 |
| Influenza A virus HA | 349 | 329 | 1.4 | 0.06 (0.06) | 0 | 0.0093 | **<0.001** | **<0.001** | 0.95 | 0.74 | 0.98 | 5 | 0 |
| HIV-1 *RT* | 476 | 335 | 6.6 | 0.086 (0.093) | 0 | 0.048 | **<0.001** | **<0.001** | 0.15 | 0.052 | 1 | 17 | 1 |

**1H** is the standard Muse-Gaut style model which only permits single nucleotides to substitute instantaneously.

**2H** is the 1H model extended to allow two nucleotides in a codon to substitute instantaneously with rate *δ* (relative to 1H synonymous rate).

**3HSI** is the 2H model extended to allow three nucleotides in a codon to substitute instantaneously if the change is synonymous (e.g., serine islands), with relative rate *Ψ*_*s*_.

**3H** is the 2H model extended to also permit any three-nucleotide substitutions, with relative rate *Ψ*.

**3H+** is the 3HSI model extended to also permit any three-nucleotide substitutions, with relative rate *Ψ*.

The nested models can be compared using standard likelihood ratio tests, using the 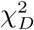 asymptotic distribution to assess significance, where *D* is chosen based on the number of constrained parameters. Key analysis results are summarized in Table S1.

#### 1. Evidence for multiple hits is pervasive

In ten of thirteen datasets the analyses strongly reject the hypothesis that 2H have zero rates, with *p<* 0.001 (2H:1H comparison). For five of thirteen datasets, we can further reject the hypothesis that 3H have zero rates (3H+:2H comparison) at *p≤* 0.05.

#### 2. Varied patterns for rate preferences

Even in this small collection of datasets, the entire spectrum of options is spanned: for the Primate Lysozyme dataset there is no evidence for anything other than 1H changes, to the Vertebrate Rhodopsin dataset, where each of the individual rates is significantly different from 0. HIV-1 *vif* dataset is the only dataset that does not support 2H rates, but does support 3H rates. Five datasets share a pattern: reject 1H in favor of 2H, and 1H in favor of 3H+, but none of the others, which can be interpreted as support for 2H rates, but none of the 3H rates.

#### 3. Varied extent of site level support for MH

Ratios between site-level likelihoods under individual models, denoted here as ER (evidence ratios), can indicate which model provides better fit to the data at a particular site. The number of sites with strong (*ER>* 5) preference for 2H vs 1H model was positive for all models rejecting 1H in favor of 2H with LRT, and ranged from 3 to 46, while a smaller number of sites (0−6) preferred 3H+ to 1H. Interestingly, for Camelid VHH, where the LRT rejects 1H in favor of 3H+, no individual sites had *ER>* 5, implying that the support for this model came from a number of individual weak site contributions.

#### 4. Interaction between 1H, 2H and 3H rates

Assuming that the biological process of evolution does include MH events, not including those in the model might have the effect of *inflating* other rate estimates. In line with other studies (Dunn *et al*., 2019), the addition of 2H rates lowers the point estimate of *ω* rates for all datasets where 2H:1H comparison is significant at *p≤* 0.05 (Table S1), sometimes dramatically (e.g., by a factor of 0.6*×* for the *β*−globin gene) which could be indicative of estimation bias due to model mis-specification. Similarly, the *δ* rate under the 2H model is always higher than the rate estimate under the 3H+ model, implying that the 2H rate may be “absorbing” some of the 3H variation. We will later see the same pattern emerge in large-scale sequence screens.

To bolster one’s intuitive understanding of model preferences, we visualized inferred substitutions at four archetypal sites in the Vertebrate Rhodopsin alignment (Yokoyama *et al*., 2008) where every single rate in the 3H+ model was significantly non-zero (Figure 1). We used joint maximum likelihood ancestral state reconstruction under the 3H+ model to estimate the number and kind of substitutions that occurred at each site (this number is a lower 6 bound and is subject to estimation uncertainty; here we use it for illustration purposes). Site 37 is what one might call a traditional single-hit substitution site, where the 1H model is preferred to all other models based on ER values; all apparent substitutions involve changes at a single nucleotide, hence the standard 1H is perfectly adequate. Of 330 codons, 149 had a preference for the 1H model compared to the 2H model. Site 144 has a dramatic preference for the 2H model over the 1H model (*ER>* 300); of 6 total substitutions, 4 involved a change at 2 nucleotides (and none – at 3). Site 281 has a preference for the 3HSI model over the 2H model (*ER* = 39), and has a complex substitution pattern : nine 1H, four 2H, and two 3H substitutions; both 3H substitutions at this site involve synonymous changes between serine codon islands (TCN and AGY). 148 other sites had a preference (*ER>* 1) for 3HSI over 2H. Finally, site 236 prefers 3H to 3HSI (*ER* = 5.4) as the only 3H substitution at that site does not involve serine.

**FIG. 1.**
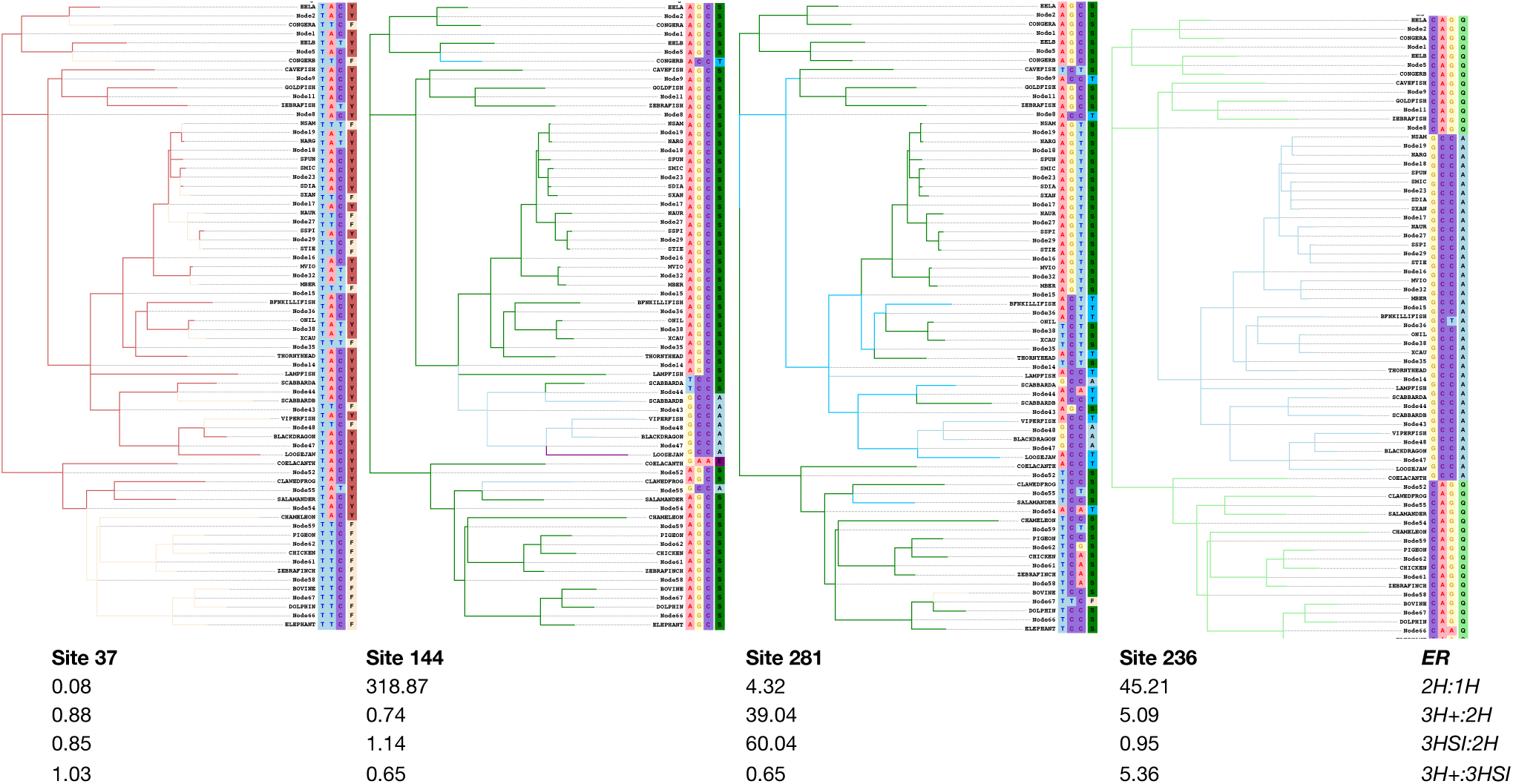
Archetypal sites based on model preferences. Four alignment sites from the Vertebrate Rhodopsin (Yokoyama *et al*., 2008) chosen illustrate substitution patterns which give rise to support for specific rate models. Branches are colored by the amino-acid that is observed/estimated to exist at the end of the branch. Internal nodes are labeled with ancestral states inferred under the 3H+ model. Evidence ratios, which are the ratios of MLE site likelihoods under the respective models, for four pairwise model comparisons are listed below each site.

**FIG. 2.**
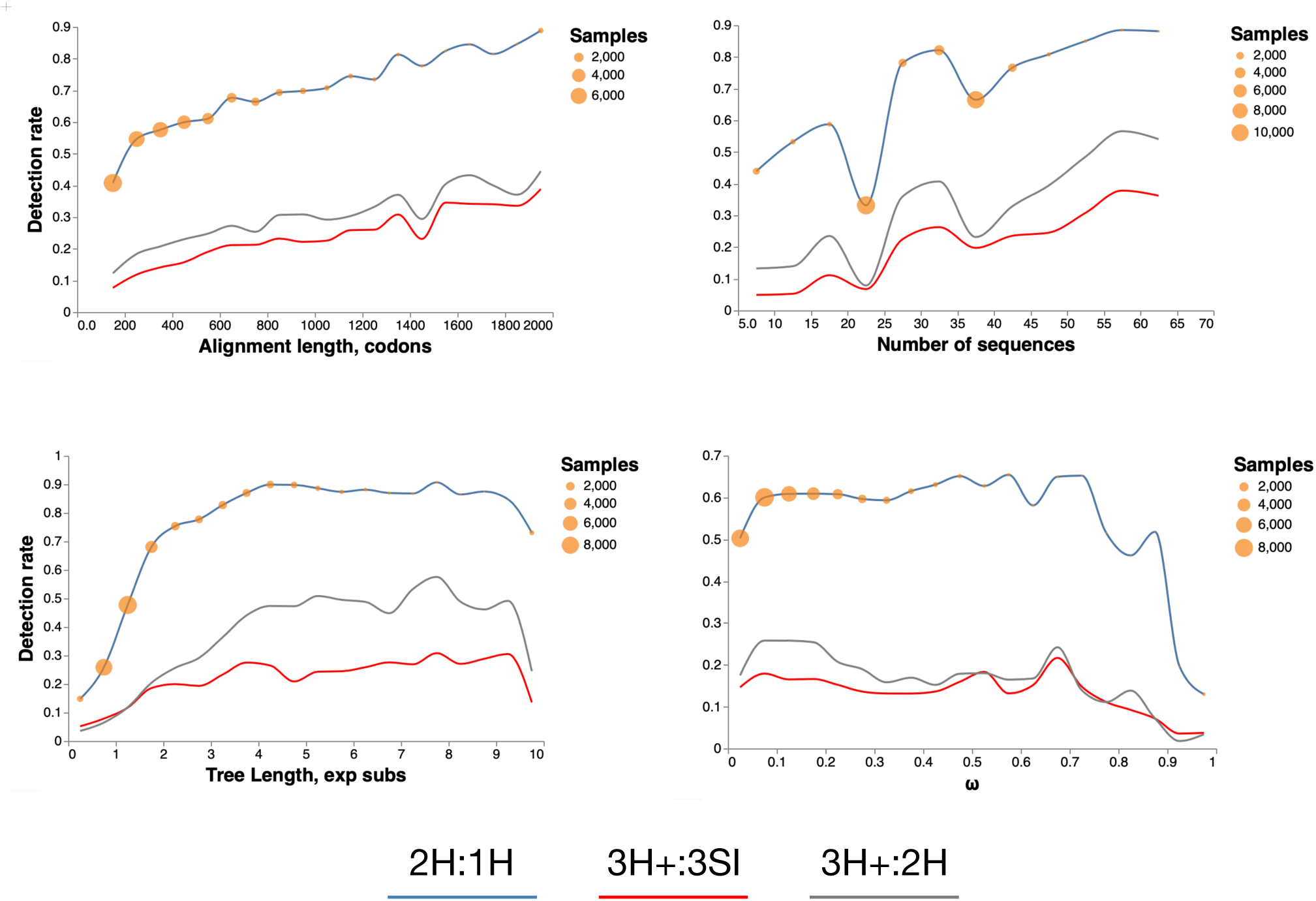
Fraction of alignments with MH rates. The fraction of alignments where the corresponding test was significant at *p ≤*0.01 as a function of alignment properties. Orange circles depict the binning steps and the number of alignments in each bin. Curves are smoothed using monotone splines. Tree lengths and *ω* values are estimated under the 1H model.

### Large-scale empirical databases

We fitted the hierarchy of MH models to 35,117 empirical datasets (Enard *et al*., 2016; Moretti *et al*., 2014; Shultz and Sackton, 2019), assembled from three large-scale studies of natural selection of nuclear genes, and a smaller collection vertebrate and invertebrate mitochondrial genes (Mannino *et al*., 2020), which represent a different evolutionary landscape (e.g., not affected by polymerase zeta).

#### Strong evidence for non-zero multiple-hit rates

We found widespread statistical support for models which includes non-zero rates involving multiple nucleotides. The 1H model was overwhelmingly rejected in favor of the 2H model (Table 3), and the improvement in fit was quite dramatic on average (mean LR), for all but the Enard *et al*. (2016) collection. A substantial fraction of alignments preferred models that allowed non-zero three rates over the 2H model, and also the 3H+ model which does not limit 3H instantaneous changes to only synonymous codons. Based on the results of the four likelihood ratio tests, each dataset could be assigned to a unique *rate preference* category Figure 3. For example, 11,899 alignments preferred 2H to 1H model, but none of the other comparisons were significant, i.e there was no evidence for non-zero 3H instantaneous rates. 2,675 alignments preferred 2H to 1H, and 3H+ to 2H, i.e. provided evidence for non-zero 3H instantaneous rates. 483 alignments preferred 2H to 1H and 3HSI to 2H, but not 3H+ to 3HSI, implying that all 3H changes were constrained to synonymous codon islands.

**Table 3.** Evidence for multiple hit rates in empirical datasets. For each collection, the fraction of alignments with significant (*p<* 0.01, based on a 5-way conservative Bonferroni correction for FWER of 5%) LRT test results, and the average value of the likelihood ratio test statistic (for significant tests) in parentheses.

|  | 2H:1H | 3H+:2H | 3H:1H | 3H+:3HSI | 3HSI:2H |
| --- | --- | --- | --- | --- | --- |
| Invertebrate mtDNA | 92% (119.2) | 7.4% (17.12) | 92% (122.2) | 8.9% (19.97) | 2.3% (9.089) |
| Vertebrate mtDNA | 54% (33.30) | 3.0% (16.60) | 50% (36.92) | 3.2% (15.11) | 0.69% (7.986) |
| Shultz and Sackton (2019) | 62% (32.39) | 20% (17.76) | 63% (39.87) | 21% (13.63) | 7.4% (12.84) |
| Moretti *et al.* (2014) | 76% (55.99) | 37% (21.82) | 77% (67.67) | 20% (13.56) | 29% (16.73) |
| Enard *et al.* (2016) | 28% (15.69) | 5.4% (14.18) | 28% (20.39) | 5.3% (10.49) | 3.4% (11.07) |
| Overall | 58% (42.91) | 22% (20.08) | 58% (52.08) | 16% (13.33) | 14% (15.70) |

**FIG. 3.**
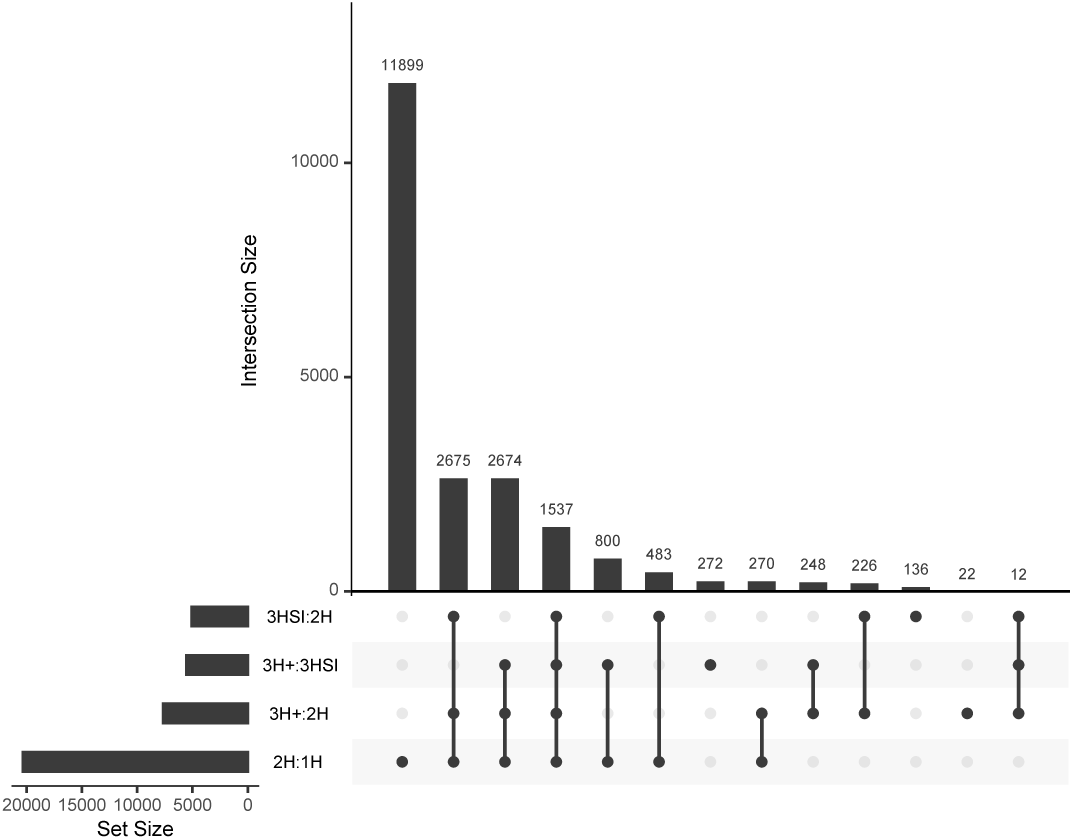
Intersections of likelihood ratio test significance. Overlaps of empirical alignments with *p≤*0.01 according to each of four LRTs performed for the combined empirical datasets. Groups of alignments for which a particular combination of tests was significant are shown in the table, with the significant tests indicated with filled dots. For example, there are 1537 alignments where all 4 tests are significant, and 136 alignments where the only significant test is 3SHI:2H.

**FIG. 4.**
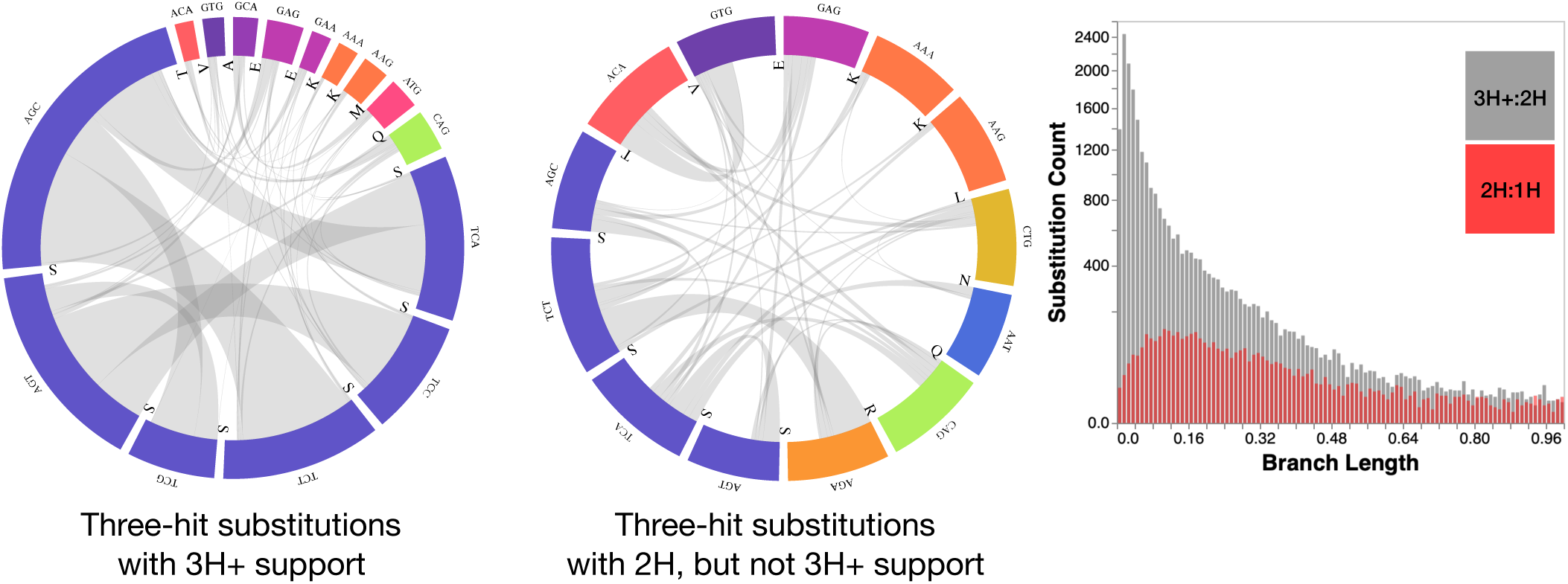
Three hit substitutions commonly occurring in empirical data. A subset of common three-hit substitutions across all empirical datasets. Three-hit substitutions with 3H+ support are defined as those occurring at sites with *ER*(3*H*+: 2*H*) *>* 5. Three-hit substitutions with 2H but not 3H+ support are defined as those occurring at sites with *ER*(3*H*+: 2*H*) *<* 1 and *ER*(2*H* : 1*H*) *>* 5. Branch lengths along which the two types of substitutions are inferred to occur are shown in the histogram.

#### Factors associated with MH detection

The rates at which 2H, 3H and 3HSI rates were detected with *p<* 0.01 as functions of simple statistics of the alignments, are shown in Figure 2. Larger (more sequences) and longer (more codons) alignments generally elicited higher detection rates for all types of multiple-hit rates. Increasing overall divergence levels between sequences, measured by the total tree length, also corresponded to increasing detection rates, up to a saturation point. The mean strength of selection, measured by the gene-average *ω* had little effect on detection rates, except for the noticeable dip for the higher values. In a simple logistic regression using 2H:1H *p<* 0.01 as the outcome variable, sequence length, and number of sequences were positively associated with the detection rate (*p<* 0.0001), while tree length was confounded with the number of sequences and was not independently predictive, and *ω* was not significantly predictive.

#### Strong MH signal comes from a small fraction of sites

For alignments where there was significant evidence for nonzero 2H and/or 3H rates (*p<* 0.01), a small fraction of sites strongly (*ER>* 5) supported the corresponding MH model. For the 2H:1H comparison, a median of 0.67% (interquartile range, IQR [0.21%−1.7%]), and for the 3H:2H comparison, a median of 0.52% (IQR [0.26%−0.94%]) (Figure S2).

#### Patterns of substitution associated with MH rates

Substitutions between serine islands (AGY and TCN) appear to be the most frequent inferred 3H change in biological alignments (see Fig **??**). Six of the most common substitutions at sites with high ER in support of the 3H+ model involve island jumping, but other amino-acid pairs are also involved in hundreds of apparent substitutions, e.g. *ATG*(*M*) *↔GCA*(*A*) Of the 7664 datasets that reject the 2H model in favor of the general 3H+ model, 2901(37.9%) fail to reject 3HSI in favor of 3H+, implying that they only require non-zero rates for synonymous island jumps. However, many of the same changes frequently appear at sites that do not strongly prefer 3H+ to 2H model, but strongly prefer 2H to 1H model (i.e, 2H sites). A key determinant of whether or not an AGY:TCN or other 3H change benefits from non-zero *Ψ* rates is the length of the branch. Branches with 3H changes that supported 3H+ model were significantly shorter than those where 2H model was sufficient: median 0.09 substitutions/site, vs median 0.26 substitutions/site. Consequently, the need to explain 3H changes happening over short branches (shorter evolutionary time, slower overall rates) provides evidence in support of 3H+ models.

Among 3H non-synonymous substitutions (see Fig 5) codons encoding for serine are still prominently represented, but not as dominant, with numerous substitutions involving methionine and other amino-acids.

**FIG. 5.**
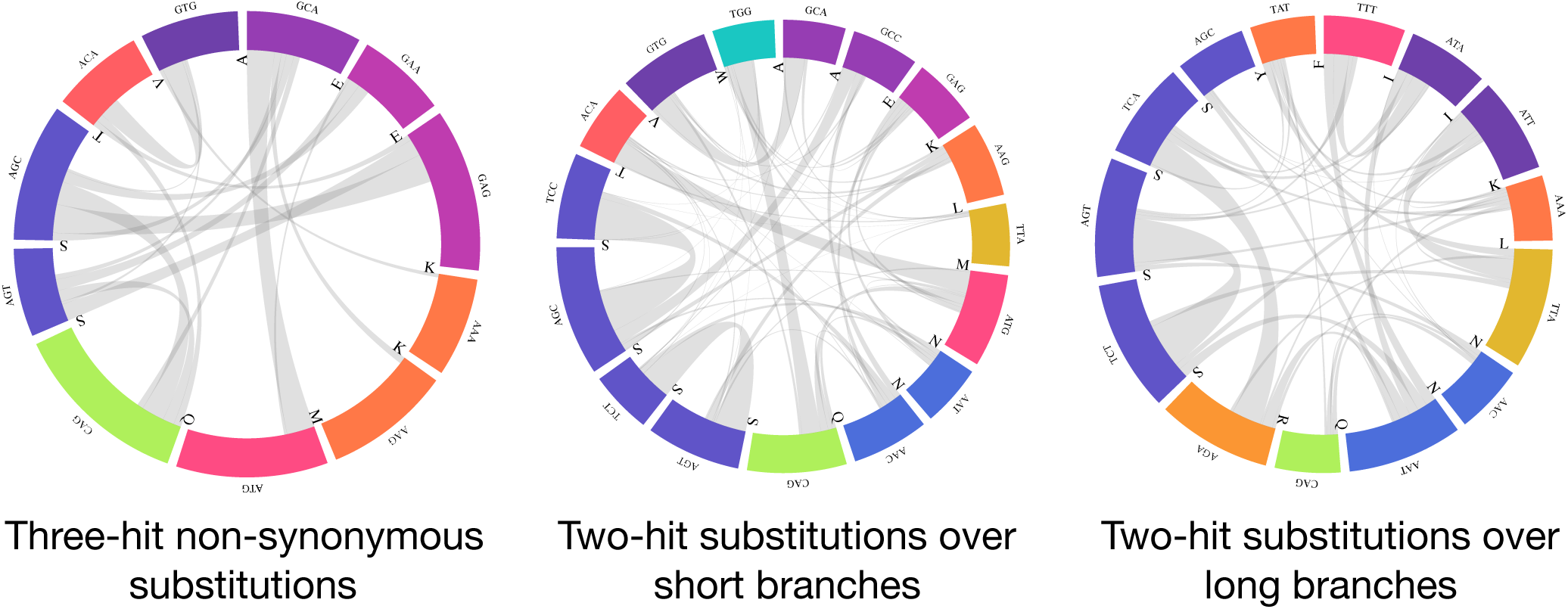
Three hit non-synonymous substitutions and two hit substitutions occurring in empirical data. A subset of common substitutions across all empirical datasets. Three-hit substitutions with non-synonymous support are defined as those occurring at sites with *ER*(3*H*+: 3*HSI*) *>* 5. Two-hit substitutions over short branches are defined as those occurring at sites with *ER*(3*H*+: 2*H*) *<* 1 and *ER*(2*H* : 1*H*) *>* 10 and branch length is *≤*0.05 subs/site. Two-hit substitutions over short branches are defined as those occurring at sites with *ER*(3*H*+: 2*H*) *<* 1 and *ER*(2*H* : 1*H*) *>* 10 and branch length is *≥* 0.25 subs/site.

Serine codons are similarly frequently involved in 2H substitutions, along both short and long branches (e.g. between codons such as *AGC ↔TCC* and *AGT ↔TCT*), but other pairs are exchanged at least 90 times, including *ACA*(*T*) *↔ATG*(*M*) and *CAG*(*Q*) *↔TGG*(*W*) (short branches) and *ATT* (*I*) *↔TTA*(*L*) (long branches).

#### Interaction between rate estimates

As with the benchmark datasets, the inclusion of multiple hit rates in models has an effect on other substitution rates. The gene wide point estimate of *ω* is systematically lowered by the inclusion of non-zero *δ* rates, even though there are rare instances when the *ω* estimates are increased (Figure S1). A Thiel-Sen robust linear regression estimate yields *ω*(2*H*) *∼* 0.965*×ω*(1*H*), but for 1150(5.7%) of the datasets with where 2H:1H comparison was signifiant, the *ω*(2*H*) *<* 0.75*×ω*(1*H*). Consequently the estimation bias in important evolutionary rates due to model mis-specification for some of the datasets could be significant. The inclusion of 3H components in the model, lowers the 2H rate even more dramatically, *δ*(3*H*+) *∼* 0.77*×δ*(2*H*).

### Simulations

#### False positive rates

We evaluated operating characteristics of the likelihood ratio tests (LRT) for MH model testing on parametrically simulated data. In the simplest case of a single-branch (two-sequence) null data generated under the 1H model, Type I error rates for 2H:1H and 3H:2H tests were on average below nominal. However, once the level of sequence divergence became very high (e.g., *>* 3 expected substitution per site), the test became somewhat anti-conservative, which is not surprising for severely saturated data (Fig. **??**). Individual branches that are this long are highly abnormal in biological datasets. Expanded to multiple sequence alignments generated using parameter estimates from four biological datasets, simulations confirmed that all the tests employed appear to be somewhat conservative; this is by design because asymptotic distributions of LRT statistics on when null hypotheses are on the boundaries of the parameter space are less conservative that the 1- or 2-degree of freedom *χ*^2^ distributions we use here (Self and Liang, 1987).

#### Power

The tests are generally well powered, especially if the effect sizes (magnitudes of MH rates) are sufficiently large (Table 4). The power to detect two-hit substitution (2H:1H) is especially high (*>* 90%) across all simulations. The test which attempts to identify non-zero triple-hit synonymous island rates (3HSI:2H) is the least powerful, because its signal is derived from a tiny fraction of all substitutions, i.e. the effective sample size is smaller that for the other tests.

**Table 4.** Power to detect MH rates. The fractions of simulations datasets that had corresponding *p<* 0.05. *N* = number of simulations in each category, and the explicit definition effect size is stated for each test.

| Test | All |  | Large effect |  |
| --- | --- | --- | --- | --- |
|  | N | Power | N | Power |
| 2H:1H | 1956 | 94% | 967 | 99%($\delta > 0.5$) |
| 3H+:2H | 1940 | 64% | 1056 | 83%($\psi > 0.5$) |
| 3HSI:2H | 447 | 33% | 114 | 51%($\psi_s > 5.0$) |
| 3H+:3HSI | 1940 | 66% | 1056 | 86%($\psi > 0.5$) |

#### False positives due to alignment errors

Whelan and Goldman (2004) suggested that non-zero estimates of triple-hit rates could be at least partially attributed to alignment errors. It is impossible, with a few rare exceptions, to declare that any particular alignment of biological sequences is correct. Hence, in order to estimate what, if any, effect potential multiple sequence alignment errors might have on our inference, we simulated one-hit data with varying indel rates with `Indelible`(Fletcher and Yang, 2009), inferred multiple sequence alignments `MAFFT` (Katoh *et al*., 2002) in a codon-aware fashion, inferred trees using neighor-joining, and performed our hierarchical model fit. This procedure induces multiple levels of model misspecification, and errors: `Indelible` uses a different model (GY94 M3) to simulate sequences, there is alignment error, and there is phylogeny inference error. Sufficiently high indel rates coupled with other inference errors can indeed bias our tests to become anti-conservative, although these levels are higher than what we see (based on per-sequence “gap”/character) ratios for our biological alignments. Empirical alignments have gap content that is consistent with alignments simulated with 0.01−0.015 indel rates, for which test performance is nominal. However, care must be taken not to over-interpret MH findings when the alignments are uncertain.

## Discussion

Nearly three in five empirical alignments considered here provide strong statistical support that at least some of the substitutions are not well modeled by standard codon substitution models that permit only single nucleotide changes to occur instantaneously. More than one in five prefer to have three-hit substitutions “enabled” by the models. Substitutions involving serine codons, which are unique among the amino-acids in that it is encoded by two codon islands which are two or three nucleotide changes from each other, are prominent in driving statistical signal for these preferences, especially if they occur along short branches. Many other amino-acid pairs are also involved in such exchanges, indicating that not all of the statistical signal is due to serine codons, although in a typical alignment only a small fraction of sites (about 1%) prefer multiple hit models strongly.

Many previous studies have provided evidence that evolutionary models that permit multiple hits provide better fit to the data or are relatively common, but the scale of this phenomenon in the comparative evolutionary context has not been fully appreciated, although the interest in model development in this area is being rekindled. Our results also show that the inclusion of multiple-hit model parameters changes *ω* estimates, and with them – potentially alter inferences of positive selection, which was demonstrated for one class of such tests by Venkat *et al*. (2018), and for data simulated with multiple hits but analyzed with standard models by Dunn *et al*. (2019).

How much of this apparent support for multiple-hits comes from biological reality, and how much from statistical artifacts, or other unmodeled evolutionary processes – the so-called phenomenological load (Jones *et al*., 2018)? Our simulation studies provide compelling evidence that the tests we use here are statistically well behaved and possess good power, i.e. our positive findings are unlikely to be the result of statistical misclassification. Other confounders, especially alignment error, have the potential to mislead the tests, but only at levels that appear higher than what is likely present in most biological alignments. In addition, there are some datasets (e.g., HIV reverse transcriptase), where alignment is not in question due to low biological insertion/deletion rates or structural information, and these data still support non-zero multiple-hit rates as well.

There is an abundance of data and examples of doublet substitutions in literature, and mechanistic explanations, e.g., due to polymerase zeta (Harris and Nielsen, 2014) exist. There are several papers arguing that the numbers of apparent triple hits occurring in sequences is greater than what we would expect solely from random mutation (Bazykin *et al*., 2004; Schrider *et al*., 2011; Smith and Hurst, 1999), however the mechanism (if it exists) by which they might occur is obscure. Sakofsky *et al*. (2014) have suggested that DNA repair mechanisms could help explain multi-nucleotide mutations, thus plausible mechanisms for triple-nucleotide changes do exist. Our analyses indicate that much, but not all, of the support for non-zero triple hit rates derives from serine codon island jumping, particularly in cases when this must occur over a short branch in the tree. Comparative species data might lack the requisite resolution to discriminate between instant multiple base changes and a rapid succession of single nucleotide changes spurred on by selection; the literature is split on which mechanism is primal (Averof *et al*., 2000; Rogozin *et al*., 2016). Such a common phenomenon is worth further investigation, in our opinion.

Our evolutionary models are broadly comparable to several others that have been published in this domain, some of which have more parametric complexity Dunn *et al*. (2019), or consider effects substitutions spanning codon boundaries Whelan and Goldman (2004). Their novel contributions are direct tests for the contributions derived from synonymous island jumping, and a simple evidence ratio approach to identify and categorize specific sites that benefit from non-zero multiple hit rates. These models are easy to fit computationally, with roughly the same cost as would be required for an *ω*−based positive selection analysis, and we provide an accessible implementation for researchers to use them. Further modeling extensions, e.g. the inclusion of synonymous rate variation, branch-site effects, etc., can be easily incorporated.

## Methods

### Substitution models

The most general model considered here is the the 3H+ substitution model and all others can be derived from it as special cases (Table 1). The model is a straightforward extension of the Muse-Gaut style time-reversible, continuous Markov processes model (Muse and Gaut, 1994). The instantaneous rate for substitutions between codons *i* and *j* (*i≠j*) is one of the six expressions defined in Table 5. *θ*_*ij*_ denote nucleotide-level biases coming from the general time reversible model (5 parameters), and *π*_*j*_ are codon-position specific nucleotide frequencies estimated from counts using the CF3×4 procedure (Kosakovsky Pond *et al*., 2010). *ω*^*k*^ are non-synonymous / synonymous rate ratios which vary from site to site using a random effect (*D*-bin general discrete distribution, *D* = 3 by default, 2*D*−1 parameters). *δ* is the rate for 2H substitutions relative to the synonymous 1H rate (baseline), *Ψ* – the relative rate for non-synonymous 3H substitutions, and *Ψ*_*s*_ – the relative rate for synonymous 3H substitutions. All parameters, except *π*, including branch lengths are fitted using directly optimized phylogenetic likelihood. Initial estimates for branch lengths and *θ* are obtained using the standard nucleotide GTR model, and models are fitted in the order of increasing complexity (1H, then 2H, then 3HSI, then 3H+), using parameter estimates from from each stage as initial guesses for the next stage.

**Table 5.** Types of modeled substitutions. Six cases for instantaneous rates *q*_*ij*_ of substituting codon *i* with codon *j* (*i* ≠ *j*). The count columns shows the number of rate matrix entires in each class (excluding the diagonal) for two commonly used genetic codes.

| Type | Expression for $q_{ij}$ | Example | Count | |
| --- | --- | --- | --- | --- |
|  |  |  | Universal | mtDNA |
| 1H synonymous | $\theta_{ij}\pi_j$ | $ACA \rightarrow ACT: \theta_{CT}\pi_T^3$ | 134 | 128 |
| 1H non-synonymous | $\omega^k \theta_{ij}\pi_j$ | $AAA \rightarrow AGA: \omega^k \theta_{AG}\pi_G^2$ | 392 | 380 |
| 2H synonymous | $\delta \prod_{n=1}^2 \theta_{ij}^n \pi_j^n$ | $CTC \rightarrow TTA: \delta \theta_{CT} \theta_{AC} \pi_T^1 \pi_A^3$ | 28 | 16 |
| 2H non-synonymous | $\delta \omega^k \prod_{n=1}^2 \theta_{ij}^n \pi_j^n$ | $AAA \rightarrow ACC: \delta \omega^k \theta_{AC} \theta_{AC} \pi_C^2 \pi_C^3$ | 1540 | 1500 |
| 3H synonymous | $\psi_s \prod_{n=1}^3 \theta_{ij}^n \pi_j^n$ | $AGC \rightarrow TCA: \psi_s \theta_{AT} \theta_{CG} \theta_{AC} \pi_T^1 \pi_C^2 \pi_A^3$ | 12 | 12 |
| 3H non-synonymous | $\psi \omega^k \prod_{n=1}^3 \theta_{ij}^n \pi_j^n$ | $GTG \rightarrow TAC: \psi \omega^k \theta_{GT} \theta_{AT} \theta_{CG} \pi_T^1 \pi_A^2 \pi_C^3$ | 1554 | 1504 |

#### Hypothesis testing

Nested models are compared using likelihood ratio tests with 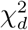 asymptotic distribution used to assess significance. *d* = 1 for 2H:1H, 3SHI:2H, and. 3H+:3HSI comparisons, *d* = 2 for 3*H*+: 2*H* comparison, and *d* = 3 for 3H+:1H comparison.

#### Comparisons to other models

The substitution model from Venkat *et al*. (2018) is very similar to our 2H model, except that *θ*_*ij*_ in their model follows the HKY85 parameterization, and it is possible to allow *κ* (transition/transversion ratio) to be different between 1H and 2H changes, and frequencies are parametrized as in the Goldman Yang model, where target codon frequencies are used in *q*_*ij*_, (Goldman and Yang, 1994).

The ECM model from Kosiol *et al*. (2007) directly estimates numerical rates for all pairs of codon exchanges in the GY94 frequency framework from a large training dataset. However, the patterns of exchangeability between codons in the ECM model captures relatively frequent exchanges between serine codons, which were further reinforced by a codon partitioning analysis of the resulting rate matrix.

The SDT model of Whelan and Goldman (2004) uses a context-averaging approach to include the effect of substitutions that span codon boundaries, and is difficult to directly relate to our models; the 3H model might be the closest to the SDT model. Regrettably, there doesn’t seem to exist a working implementation of the SDT model (pers. comm from Simon Whelan), which makes direct comparison to our approaches impractical.

The KCM model of (Zaheri *et al*., 2014) only has a single rate for multiple hits (double or triple), and has position-specific nucleotide substitution rates (*θ* in our notation), so it would be most comparable to the 3H model with *δ* = *Ψ*.

The GPP model class of Dunn *et al*. (2019) can be parametrized to recapitulate our models because it can capture (in a log-linear parametric form), arbitrary rate matrices with suitable parametric complexity. Several of the models considered in Dunn *et al*. (2019) include multiple hits, but they are not directly comparable to ours, mostly because they also incorporate *ω* rates that depend on physicochemical properties of amino acids, and because the exact parametric form of the models are hard to glean from available description.

#### Empirical data

The Moretti et al (Selectome) data collection consists of 13,303 gene alignments from the *Euteleostomi* clade of Bony Vertebrates from Version 6 of the database (Moretti *et al*., 2014) and can be downloaded from data.hyphy.org/web/busteds/.

The Shultz et al data collection (Shultz and Sackton, 2019) contains 11,262 orthologous protein coding genes from 39 different species of birds and is freely available at https://datadryad.org/stash/dataset/doi:10.5061/dryad.kt24554.

The Enard et al data collection (Enard *et al*., 2016) includes 9,861 orthologous coding sequence alignments of 24 mammalian species and is available at https://datadryad.org/stash/dataset/doi:10.5061/dryad.fs756.

Our mtDNA data set consists of both invertebrate and vertebrate Metazoan orders with gene alignments of each of the 13 protein coding mitochondrial genes. This data set was compiled from NCBI’s GenBank for Mannino *et al*. (2020) and can be found at: https://github.com/srwis/variancebound.

#### Simulated data

The two-sequence simulated data set was generated in HyPhy using the `SimulateMG94` package from https://github.com/veg/hyphy-analyses/. These sequences were simulated under the 1H (no site-to-site rate variation) with varying sequence and branch length as well as varied but constant *ω* across sites but no multiple hits. We generated 1000 simulations scenarios and drew 5 replicates per scenario; *ω* was drawn from *U* (0.01,2.0), branch length was drawn *Exp*(*U* (0.01,1.0)), and codon lengths as an integer from 100 to 50000 uniformly. Parameter values were sampled using the Latin Hypercube approach to improve parameter space coverage.

Multiple sequence simulations were based on the fits to one of four benchmark datasets: Drosophila adh, Hepatitis D antigen, HIV vif, and the Vertebrate rhodopsin data. We took all model parameter estimates under the 3H+ model as the starting point, and generated 500 replicates per dataset of which 35% were null (1H), 10% each from 2H, 3SHI or restricted 3H+ (*Ψ*_*s*_ = 0), and 35% from 3H+. *δ, Ψ* and *Ψ*_*s*_ parameters, when allowed to be non zero by the model, were sampled from *U* (0,1), *U* (0,1), and *U* (0,10), respectively.

Sequences with indel rate variation were generated using INDELible v1.03 (Fletcher and Yang, 2009). Indel rates were varied 0.01 to 0.06 in increments of 0.005 (100 replicates per value), and the site-to-site rate variation was modeled with a 3-bin M3 model.

#### Data availability

All of the sequence alignments, simulated or biological, and simulation/configuration scripts are available for download from data.hyphy.org/web/multihit/

### Implementation

All analyses were performed in HyPhy version 2.5.1 or later. The method used to fit the standard 1H model along with 2H, 3H and 3HSI versions is available from as the FitMultiModel package available from: https://github.com/veg/hyphy-analyses/, and can be invoked with hyphy fmm in version 2.5.7 or later.

Interactive results can be viewed at http://vision.hyphy.org/multihit using JSON results output by HyPhy.

## Acknowledgments

This work was supported by National Institutes of Health grants R01 GM093939 and R01 AI134384 to SKP.

## Supplementary Material

**Table S1.** Estimated *ω* rate distributions for benchmark datasets for different models on the benchmark datasets. *E*[*ω*] : the mean *ω* value for the 1H model.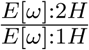: the ratio of mean *ω* estimates from 2H and 1H models. 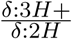: the ratio of *δ* estimates from 3H+ and 2H models. The datasets are sorted by increasing values of the 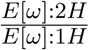 column. Genes where there was significant evidence (LRT *p<* 0.05) of non-zero 2H rates are bolded, and those where there is evidence of non-zero 3H rates is underlined.

| Gene | 1H |  |  |  |  | 2H |  |  |  | 3H+ |  |  |
| --- | --- | --- | --- | --- | --- | --- | --- | --- | --- | --- | --- | --- |
| | $E[\omega]$ | $\omega_1(p_1)$ | $\omega_2(p_2)$ | $\omega_3(p_3)$ | $\frac{E[\omega]:2H}{E[\omega]:1H}$ | $\omega_1(p_1)$ | $\omega_2(p_2)$ | $\omega_3(p_3)$ | $\frac{\delta:3H+}{\delta:2H}$ | $\omega_1(p_1)$ | $\omega_2(p_2)$ | $\omega_3(p_3)$ |
| <u>β-globin</u> | 0.28 | 0.0073 (32.4%) | 0.28 (58.9%) | 1.3 (8.66%) | 0.6 | 0.00071 (28.4%) | 0.17 (61.5%) | 0.64 (10.1%) | 0.86 | 0.0035 (30.2%) | 0.19 (61.1%) | 0.7 (8.68%) |
| **Vertebrate Rhodopsin** | 0.12 | 0.0097 (58.8%) | 0.17 (30.3%) | 0.54 (10.9%) | 0.7 | 0.0085 (60.6%) | 0.12 (29.6%) | 0.4 (9.79%) | 0.75 | 0.0088 (60.6%) | 0.13 (29.8%) | 0.41 (9.63%) |
| Flavivirus NS5 | 0.047 | 0.0026 (63.9%) | 0.07 (26.5%) | 0.28 (9.57%) | 0.73 | 0.0031 (67.1%) | 0.062 (25.5%) | 0.22 (7.4%) | 0.68 | 0.0031 (66.8%) | 0.063 (25.7%) | 0.22 (7.55%) |
| **Drosophila adh** | 0.1 | 0 (50.1%) | 0.1 (33.8%) | 0.43 (16%) | 0.73 | 0 (38.1%) | 0.047 (42.5%) | 0.29 (19.4%) | 0.76 | 0 (37.2%) | 0.046 (43.2%) | 0.29 (19.6%) |
| **COXI** | 0.07 | 0.0027 (75.4%) | 0.033 (17.3%) | 0.14 (7.33%) | 0.79 | 0.0031 (79%) | 0.035 (14%) | 0.12 (6.99%) | 1 | 0.0031 (79.3%) | 0.036 (13.9%) | 0.12 (6.87%) |
| **Sperm lysin** | 1.1 | 0.11 (36.9%) | 1.1 (41.7%) | 2.9 (21.4%) | 0.83 | 0.096 (37.2%) | 1 (43.8%) | 2.5 (19%) | 0.87 | 0.1 (37.6%) | 1 (43.7%) | 2.4 (18.7%) |
| **Hepatitis D virus antigen** | 0.48 | 0.033 (46.2%) | 0.4 (32.4%) | 1.6 (21.4%) | 0.84 | 0.038 (48.7%) | 0.37 (30.2%) | 1.3 (21.2%) | 0.9 | 0.037 (48.3%) | 0.36 (29.9%) | 1.3 (21.7%) |
| <u>Camelid VHH</u> | 0.95 | 0.12 (34%) | 0.73 (40.8%) | 2.5 (25.1%) | 0.89 | 0.1 (34.7%) | 0.66 (40.4%) | 2.2 (24.9%) | 0.92 | 0.1 (34.4%) | 0.66 (40.2%) | 2.2 (25.5%) |
| **HIV-1 RT** | 0.19 | 0.016 (71.4%) | 0.35 (22.2%) | 1.6 (6.38%) | 0.93 | 0.015 (70.9%) | 0.31 (22.3%) | 1.5 (6.82%) | 0.93 | 0.015 (71.1%) | 0.31 (22.2%) | 1.5 (6.74%) |
| Encephalitis env | 0.054 | 0.024 (77.9%) | 0.028 (15.5%) | 0.48 (6.62%) | 0.96 | 0.023 (71.8%) | 0.026 (21.6%) | 0.46 (6.64%) | 1 | 0.023 (75.8%) | 0.026 (17.6%) | 0.46 (6.64%) |
| **Influenza A virus HA** | 0.47 | 0.095 (72%) | 0.93 (23%) | 3.7 (5.06%) | 0.98 | 0.097 (72.8%) | 0.95 (22.6%) | 3.7 (4.69%) | 0.99 | 0.098 (73.2%) | 0.98 (22.4%) | 3.8 (4.41%) |
| <u>HIV-1 vif</u> | 0.84 | 0.14 (61.5%) | 0.83 (20.6%) | 3.2 (17.9%) | 0.99 | 0.16 (66.2%) | 0.96 (16.4%) | 3.2 (17.4%) | 0.16 | 0.16 (65.6%) | 0.9 (16.2%) | 3.2 (18.2%) |
| Primate Lysozyme | 0.61 | 0.14 (0%) | 0.22 (82%) | 2.4 (18%) | 1 | 0.14 (0%) | 0.22 (82%) | 2.4 (18%) | 1 | 0.21 (16%) | 0.22 (66%) | 2.4 (18%) |

**FIG. S1.**
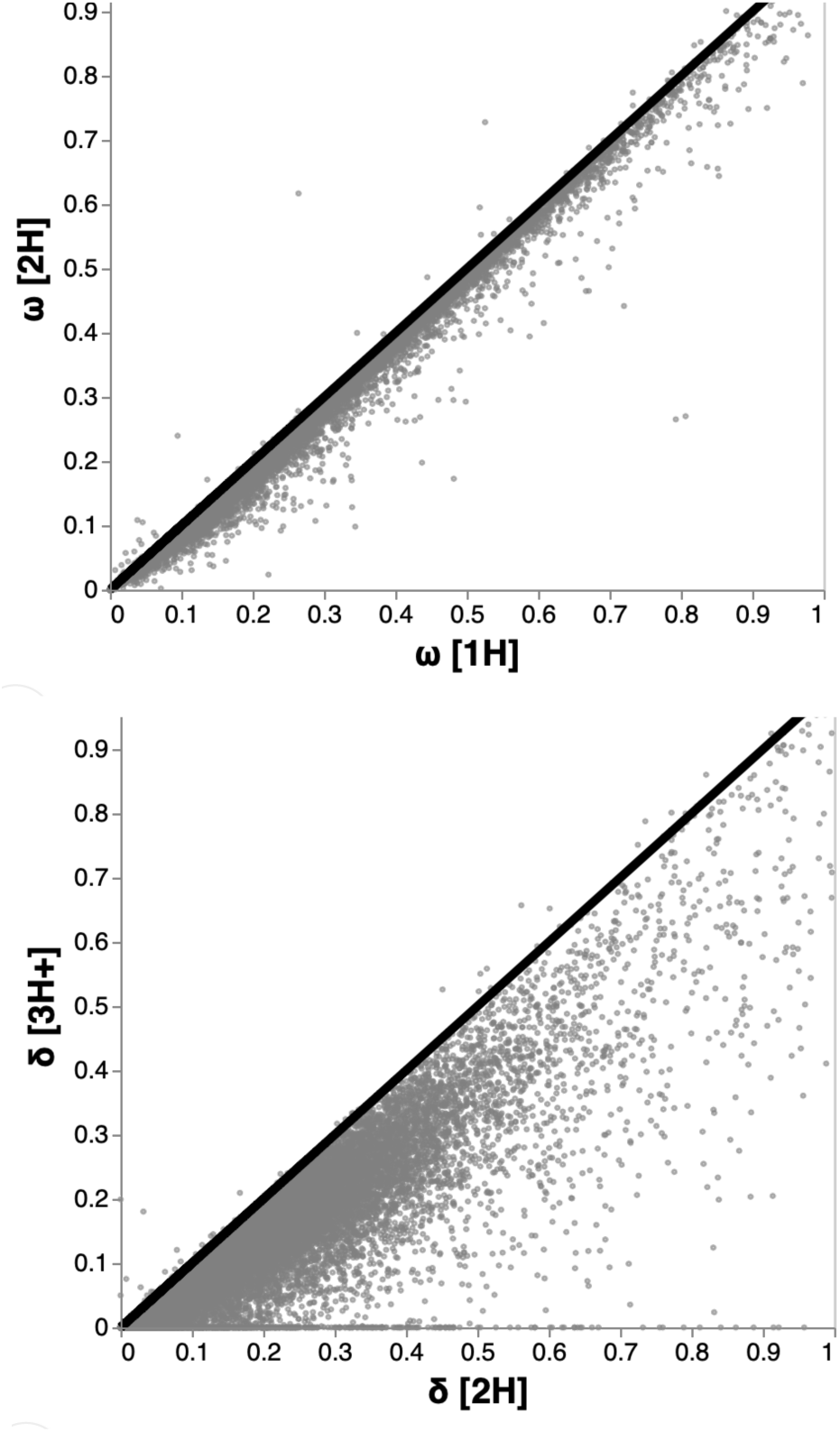
The effect of model choice on rate estimates. Point estimates of global rate parameters under different models for each of the empirical datasets.

**FIG. S2.**
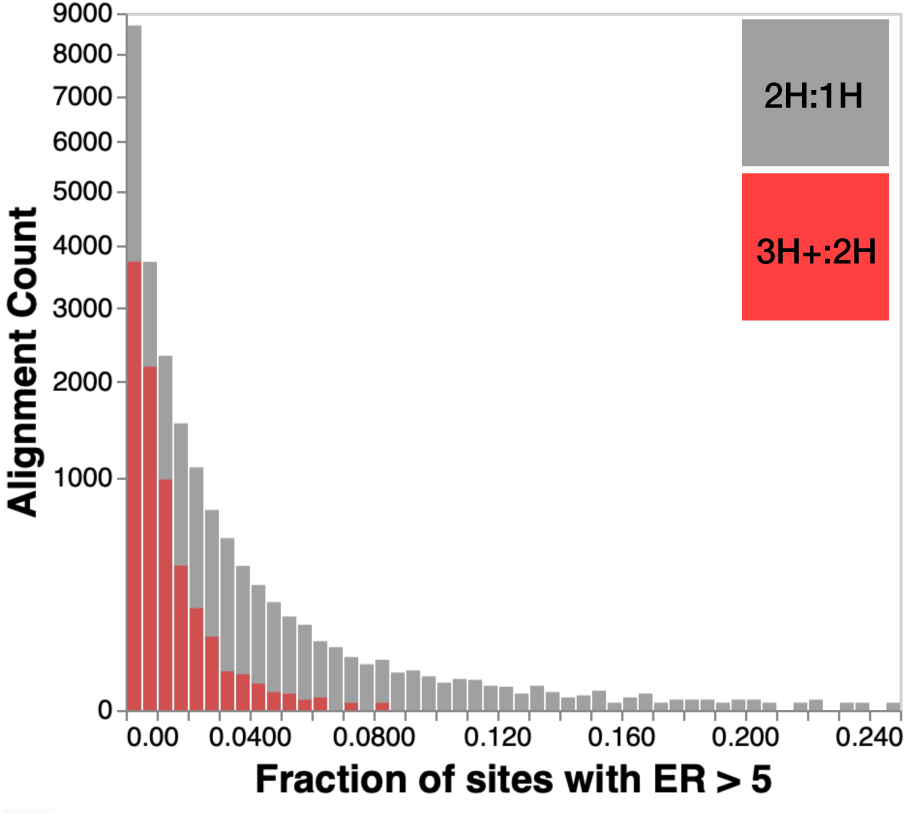
The fraction of sites with strong MH model preference. Histograms are over alignments where there was significant (*p* < 0.01) support for the corresponding model: 20,338 for 2H:1H (gray) and 7664 for 3H:2H (red).

**FIG. S3.**
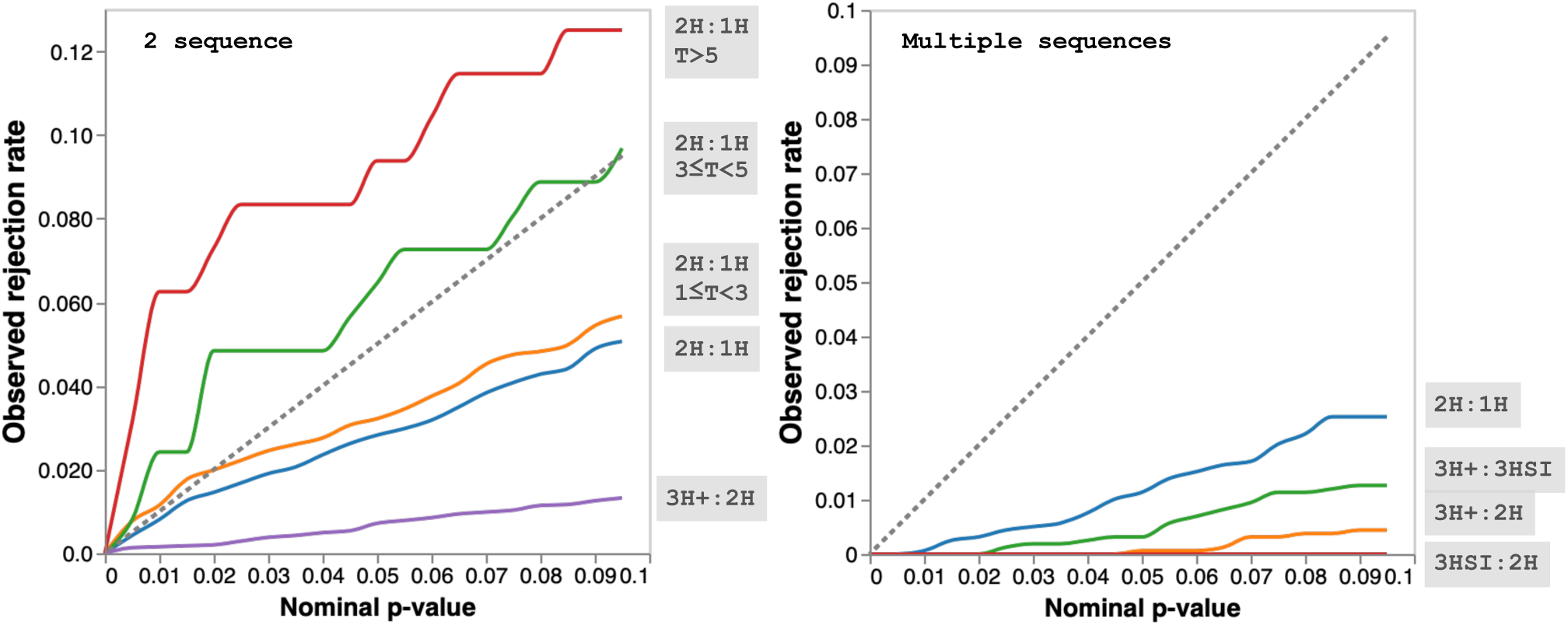
False positive rates for LRTs on simulated data. For the two sequence simulations, we further stratified the simulations by the length of the branch, *T*, measured in expected substitutions per site.

**FIG. S4.**
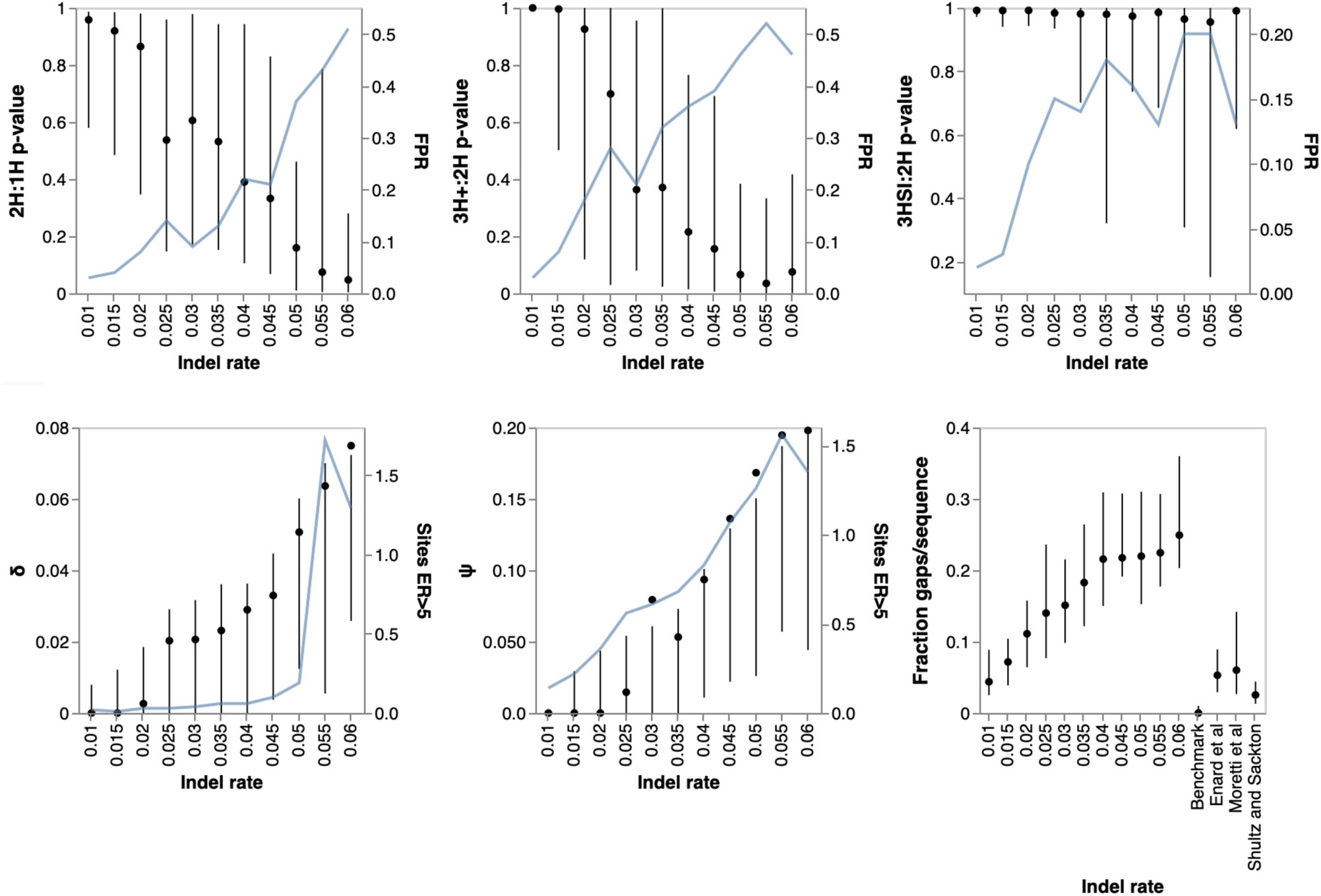
Indel Rate verse TH Rate. Alignments with indel were simulated using INDELible across using the Dropsophila *a*dh tree and alignment length using GY94 M3 model with site-to-site *ω* variation. LRT p-values and rejection rates (FPR, at *p* ≤ 0.05) are shown for different tests in the top row. The bottom row sows estimated *δ* and *Ψ* rates as a function of simulated indel rates, as well as the number of sites inferred to have high evidence ratios (ER) for 2H or 3H modes. The plot on the bottom right shows the average fraction of a sequence that in an alignment that is comprised of gaps is shown for simulated data, and empirical collections.

